# Intrinsic diving reflex enhances cognitive performance by alleviating microvascular dysfunction in vascular cognitive impairment

**DOI:** 10.1101/2024.04.25.591162

**Authors:** Willians Tambo, Keren Powell, Steven Wadolowski, Daniel Sciubba, Michael Brines, Chunyan Li

## Abstract

Vascular cognitive impairment (VCI) stands as the second-most prominent contributor to cognitive decline, lacking efficacious interventions. Chronic cerebral hypoperfusion (CCH) triggers microvascular dysfunction, which plays a critical role in VCI pathophysiology, emerging as a pivotal therapeutic target. While interventions addressing facets of microvascular dysfunction like angiogenesis and blood-brain barrier functionality show promise, the evaluation of microvascular constriction, another key component, remains unexplored. The diving reflex (DR) represents an oxygen-conserving response, characterized by robust vasodilation and potentially also inducing angiogenesis. In this investigation, we studied DR’s functionality and underlying mechanisms within a rat bilateral common carotid artery occlusion induced CCH model. Remarkably, progressive hippocampal microvascular constriction exhibited strong correlations with short-term memory impairment during both early (R^2^=0.641) and late phases (R^2^=0.721) of CCH. Implementation of DR led to a significant reduction in microvascular constriction within the hippocampus (∼2.8-fold) and striatum (∼1.5-fold), accompanied by enhanced vasodilatory capacity and heightened expression of vasoactive neuropeptides. Furthermore, DR attenuated microvascular degeneration across various brain subregions affected by CCH, concomitant with increased levels of multiple angiogenic factors. The reinforced microvascular integrity facilitated by DR corresponded with significantly improved short-term recognition memory and long-term spatial memory functions observed during the late phase of CCH. The comprehensive and synergistic effects of DR on various aspects of microvascular function and cognitive preservation highlight its potential as a disease-modifying therapeutic strategy in VCI.

## Introduction

Vascular cognitive impairment (VCI) ranks as the second most prevalent cause of cognitive decline, yet effective therapeutic strategies are lacking [1–5]. Chronic cerebral hypoperfusion (CCH), the primary underlying factor in VCI pathogenesis, initiates a cascade of vascular anomalies, including vessel constriction, microvascular depletion, and endothelial distress [2,6–8]. These pathological alterations predominantly impact brain regions essential for cognition, such as the hippocampus [9] and white matter [10], rendering them susceptible to hypoxic/ischemic insults. Despite the absence of efficacious clinical interventions, experimental approaches targeting microvascular damage, notably angiogenesis and reinforcement of the blood-brain barrier (BBB), have exhibited promise in addressing CCH [11–13]. Emerging evidence, however, underscores the significance of microvascular constriction [14–17], a pivotal aspect of microvascular dysfunction, in the progression of VCI. Consequently, we propose that interventions aimed at enhancing microvascular dilation, alongside promoting angiogenesis, may offer substantial therapeutic potential in the management of VCI.

The diving reflex (DR) represents a potent intrinsic mechanism geared towards prolonging survival amidst the hypoxic or anoxic environments experienced underwater. Its induction involves the concurrent stimulation of trigeminal sensory afferents and the onset of apnea, thereby triggering a sequence of physiological adaptations aimed at enhancing cerebral perfusion [18,19]. Notably, DR enhances cerebral perfusion through potent and targeted cerebral vasodilation [20,21]. This effect is discernible not only within the cortical regions but also extends to subcortical areas such as the hippocampus and thalamus [20], which are distinguished by a prevalence of microvasculature over large vessels [22–24]. Additionally, prior investigations have established that DR elicits pronounced peripheral vasoconstriction, facilitating the redistribution of peripheral blood flow to vital organs (e.g., brain) [19,25], a phenomenon observed in humans [26,27], rodents [20,28], and seals [29]. Of note, peripheral constriction observed in remote limb ischemic conditioning (RLIC) has been associated with the promotion of angiogenesis [30,31], suggesting a potential role for DR-induced peripheral vasoconstriction in angiogenic processes. Moreover, recent findings from our research indicate that DR triggers the activation of nuclear factor-erythroid-2-related factor 2 (Nrf2) within the rat brain [32], a crucial regulator implicated in endothelial function, neurovascular coupling, and BBB preservation in VCI [33]. Although DR shows promise in preserving various aspects of microvascular function, its effectiveness in mitigating microvascular injury and enhancing cognitive function in CCH has not been explored, to date.

To investigate the protective capabilities of DR against microvascular injury in VCI, we conducted a comprehensive study examining both structural and functional outcomes, as well as potential underlying mechanisms. This investigation utilized a well-established rat model of CCH induced by bilateral common carotid artery occlusion (2VO). To thoroughly elucidate the involvement of DR in CCH, we first characterized the temporal progression of microvascular dysfunction in the 2VO model, encompassing both constriction and degeneration, and correlated these findings with the gradual onset of cognitive decline. Subsequently, we applied a validated DR protocol [20,34–36] to animals undergoing 2VO and evaluated the effects of DR on vasodilatory and angiogenic reactions, as well as its influence on both short-term and long-term memory capabilities. To delineate the potential mechanisms underlying DR, we introduced an additional control group involving swimming. The results of this investigation suggest that DR holds considerable promise as a disease-modifying therapeutic approach for addressing VCI.

## Materials and Methods

### Experimental animals

Two cohorts of male Sprague-Dawley rats (Charles River Laboratories, USA) were used for this study. The first cohort was comprised of 48 rats (200-250g) for the initial assessment of 2VO pathophysiological development. The second cohort consisted of 48 initially juvenile (100-125g) rats for the comparative 2VO, 2VO+DR, and 2VO+swim assessments. 24 rats underwent either DR or swim training into adulthood (200-250g), while 24 rats from the same cohort aged alongside them for sham and 2VO-vehicle. For the sake of parity, all animals undergoing 2VO were induced at the same age. Experiments were performed in accordance with the *Guide for the Care and Use of Laboratory Animals* [37] following approval by the Institutional Animal Care and Use Committee.

### Experimental design

A schematic of the experimental protocols is depicted in **Fig. 1**. Rats were randomly divided into four groups: sham (N=24), 2VO (N=48), 2VO+DR (N=12), and 2VO+swim (N=12). Sham animals were comprised of animals from 2 cohorts, each comprising 12 animals. The 2VO group was further separated into 4 subgroups, 2wks (N=12), 4wks (N=12), and 6wks survival (N=12 x 2 cohorts). A power calculation was used (α=0.05), which indicated a minimum group size of N=6, for both fresh and transcardial perfusion collections. Behavioral tests were assigned an N=9, due to increased inter-animal variability; behavioral data was collected serially from all 2-VO alone animals every 2wks, leading to N=21 for 2wks and N=15 for 4wks. Every two weeks, a set of animals was collected, leading to serially decreasing N-numbers. At 2wks, 4wks, and 6wks, 2VO animals were assessed for cognitive dysfunction. At 6wks, the 2VO+DR and 2VO+swim groups underwent the same set of evaluations. At the appropriate timepoints, animals were sacrificed for histological and biochemical analyses after a final round of cognitive assessments.

**Figure 1.**
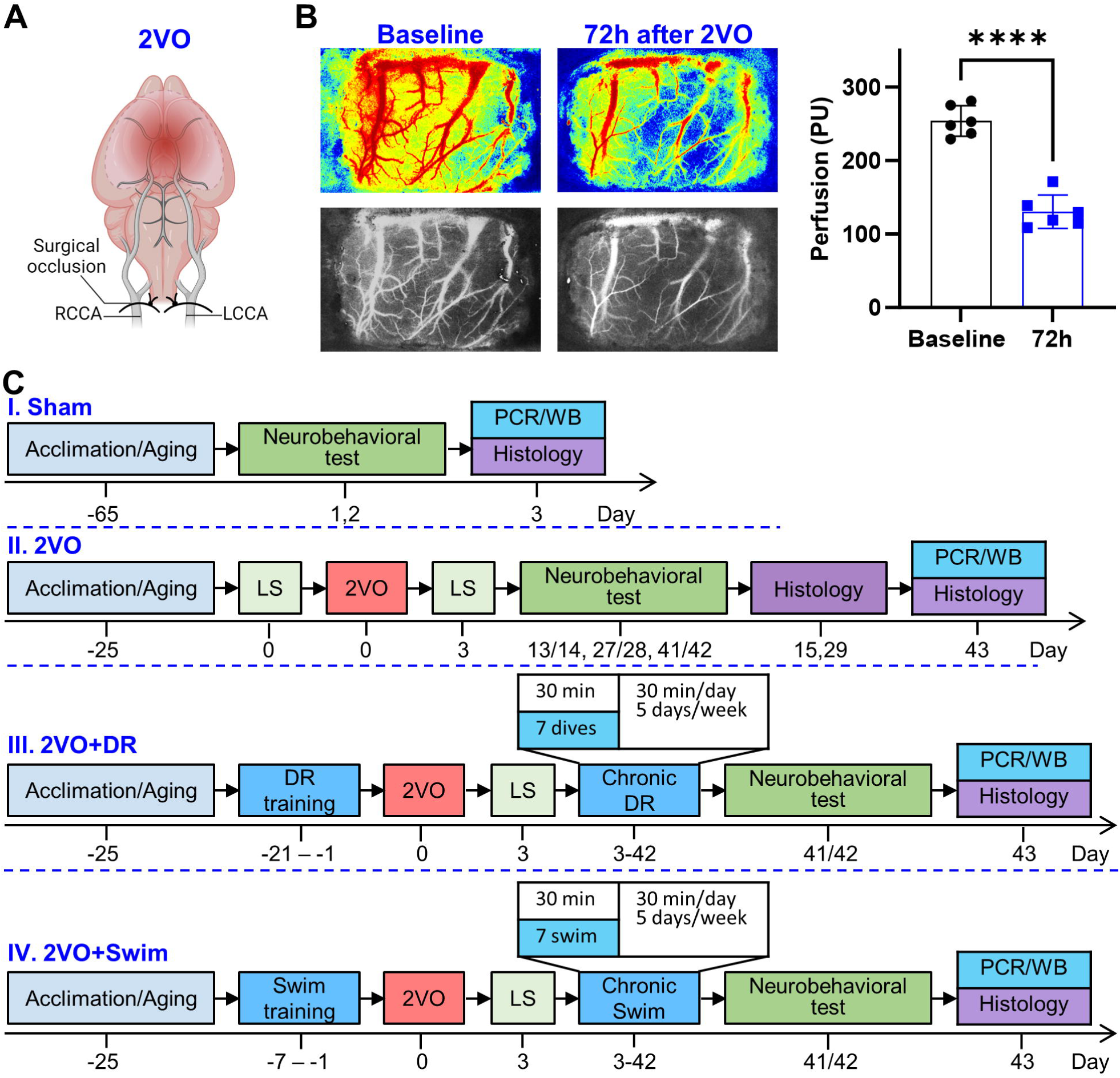
Conceptual diagram of experimental protocols. A) Schematic representation of 2VO as a rat model of chronic cerebral hypoperfusion. B) Assessment of CBF by Laser Speckle imaging indicates that 72-hours after 2VO induction, perfusion in the cortex decreases ∼50% below baseline. C) Timeline and experimental design of experimental protocols. (Abbreviations: 2VO: bilateral common carotid artery occlusion; DR: diving reflex; LS: Laser Speckle; NOR: novel object recognition test; PCR: Polymerase chain reaction; WB: Western blot)

### Rat model of chronic cerebral hypoperfusion

Male Sprague-Dawley rats were subjected to CCH via 2VO (**Fig. 1A)** [38]. Briefly, rats were anesthetized with isoflurane and placed on a thermal plate to maintain body temperature 37°C. A midline incision was made in the animal’s neck, and the left common carotid artery was exposed, gently separated from the vagus nerve, and permanently cauterized. The same procedure was followed for the right common carotid artery. Incisions were closed with non-absorbable nylon sutures (Ethicon, USA), subcutaneous buprenorphine (Covetrus, USA) was applied, and animals were returned to their home cages and observed for complications due to the surgical procedures. The sutures were removed 10-days following surgery. During the survival period, animals were monitored daily.

### Diving and swimming training

A modified version of the protocol established by McCulloch [34] was used to establish a model of rodent DR. This protocol has been confirmed for studying the mammalian diving response in mice and rats [20,35,36]. Rats underwent training in a MazeEngineers (USA) rectangular acrylic tank (100 x 38 x 15 cm) featuring a ¾-inch thick bottom and a ½-inch thick top. The tank was partitioned into three channels, each ∼100 cm in length, with starting and ending areas totaling ∼50 cm, for a total diving length of ∼2.5m. During training and DR/swim treatment, water temperature was maintained at 30±2°C.

Over the course of three weeks (5 days/week), rats underwent gradual training to swim and dive the length of the maze (Fig. 1C). The initial week focused on swimming, allowing rats to first freely acclimate. Those observed swimming were then trained to navigate the maze. The subsequent two weeks involved diving training (+0.25 meters/day). During swimming and diving training, rats underwent 3-5 trials/day. Rats were given time at the start area to encourage voluntary diving, and then time at the end platform to groom, to associate it with safety. Following the original protocol, no treats were provided to avoid activating the reward circuit [34].

### Diving and swimming treatment protocol

Starting three days post-2VO, rats underwent 30-minute sessions comprising 7 repeated diving bouts, resembling protocols employed in human assessments of DR involving multiple apneic exposures [39]. Animals in the 2VO+swim group underwent similar 30-min sessions, comprising 7 bouts of repeated swimming, for the same distance and duration as the diving animals (**Fig. 1C)**. The experimental protocol included a 5-minute resting period between each bout. During this interval, rats were transferred from the finish area to a drying area lined with clean towels, allowing them to groom and relax before the next bout. Diving/swimming sessions were repeated daily, 5-days per week, for a total of 6wks.

### Memory assessments

Behavioral assessments were performed in a dedicated assessment suite, with consistent lighting and minimal extraneous stimuli. Tests followed previously established protocols and were performed in order of least stressful to most stressful, to reduce outside interference. All assessments were recorded with the EthoVision software (Noldus, USA). After each assessment, testing objects and fields were thoroughly wiped with 70% alcohol and Peroxiguard to eliminate feces and other odors, preventing these from affecting the behavior of the other rats. Assessments were performed by a dedicated technician blinded to experimental groups.

### Novel object recognition test

The novel object recognition test (NOR) assesses the working recognition memory of rats based on the assumption that a healthy rat will interact more with a novel object rather than one to which they are acclimated [40]. The experiment was conducted in a 60x60x30 cm open field, comprised of opaque black acrylic. The NOR test consists of two phases: habituation and testing. During the habituation phase, the rats are familiarized with an object before the test. In the test phase, a novel object is introduced alongside the familiar object, and the time the rats interact with the familiar and novel objects is recorded. The two objects are of similar size and material, to increase the reliability of the test. The discrimination index (DI) is calculated as TN-TF/TN+TF (TN: time of novel object exploration; TF: time of familiar object exploration), for a possible range of: −1.0–1.0.

### Y-maze test

The Y-maze test assesses long-term spatial memory [40]. The maze is comprised of a Y-shaped structure (MazeEngineers, USA), with three blue, opaque plastic arms that are positioned 120 degrees apart from each other. Long-term spatial memory is assessed via placing the rats in the maze with one arm closed off for a period of 10-minutes. After an inter-trial interval of 4-hours, the rats are returned to the maze with the blocked arm accessible. Rats with intact long-term spatial memory will preferentially explore the arm which had previously been blocked off (novel arm). The frequency of entries into the novel arm is recorded and used to assess long-term spatial memory.

### Cerebral blood flow by laser speckle contrast image

Cerebral blood flow (CBF) was imaged using the high-resolution Laser Speckle Contrast Imaging System (RFLSI II, RWD, China) [41]. Anesthetized rats were positioned in a stereotaxic apparatus, and craniotomy was performed to remove a flap of the skull and expose the left cortex. The surface of the cortex was illuminated with a 784-nm laser (60 mW), and the blood flow was recorded for 10 s (2.8×, 2048LJ×LJ2048, 10 frames at 10 Hz with 2 ms exposure time). Rats were thermostatically controlled (37°C) on a warming plate to mitigate the impact of body temperature during CBF measurement.

### Histological specimen collection and preparation

Rats were deeply anesthetized with isoflurane and transcardially perfused with cold phosphate-buffered saline (PBS, Sigma, USA), followed by cold 4% paraformaldehyde (PFA, Sigma, USA) in PBS as a fixative. The brains were removed and immersed in PFA overnight, preserved in gradient sucrose solutions, cryo-embedded in a 1:3 mixture of 30% sucrose and Optimal Cutting Temperature Compound (Thermo Fisher Scientific, USA), and stored at −80°C for future cryosectioning. Brains were serially sectioned in a coronal orientation at 800 µm intervals using a cryostat (Leica Biosystems, Germany). 18 µm sections were mounted on Superfrost Plus glass slides (Thermo Fisher Scientific, USA) and 14 µm sections were mounted on Polysine glass slides (Thermo Fisher Scientific, USA). Slides were stored at −30°C for future usage.

### Cellular Health Evaluation using ImageJ

Samples on Superfrost slides were stained with H&E. Cellular health was assessed across brain sections using existing ImageJ software. Whole brain section images were taken using EVOS 7000 (Thermo Fisher Scientific, USA) with a 20x objective and stitched together using ImageJ Grid/Collection Stitching Tool [42] at a pixel resolution of 0.50292µm/pixel. Images were re-segmented into 515x515µm (1024x1024pixel) images and run through the following basic workflow made using ImageJ macro language. 1) Hematoxylin-stained cells were segmented using StarDist [43]. 2) Individual cells were determined to be healthy pyramidal cells, pyknotic cells, or unknown based off their morphology. 3) Healthy cells were marked green, while all other cells remained unmarked. ROIs were selected for each brain region (cortex, hippocampus CA1, amygdala, thalamus, striatum) and the total number of healthy cells was counted using the markers and Particle Analyzer Tool. Cell counts were reported as absolute number per .25 mm^2^.

### Quantification of macro- and micro-vasoconstriction

To assess macro-and micro-circulatory changes [40,44], H&E-stained slides were imaged using the EVOS M7000 (Thermo Fisher Scientific, USA) at 20x magnification. Left internal cerebral artery (ICA), middle cerebral artery (MCA), anterior cerebral artery (ACA), basilar artery (BA), and pial arterioles atop the cortex were imaged. Vessel wall thickness was measured at three-equally spaced points along the circumference of major vessels and pial arterioles and averaged. Pial arteriole diameter was also measured and a thickness to diameter ratio was calculated, to account for the baseline variations in arteriole size. Parenchymal arteriole collapse in the cortex, hippocampus CA1 and dentate gyrus (DG), amygdala, thalamus, and striatum, was identified in the whole-brain H&E images, counted manually, and presented as percentage of total arteriole number.

### Immunofluorescent labeling

Sections mounted on Polysine glass slides were subjected to FITC-lectin staining [45]. Slides were washed with tris-buffered saline plus Tween-20 (TBS-T, Sigma, USA), blocked with goat serum (Abcam, USA) supplemented with 1% bovine serum albumin (Sigma, USA) for 1 hour at room temperature, and incubated with primary antibody FITC-lectin at a dilution factor of 1:75 at 4°C overnight. Slides were counterstained with DAPI (1:2000, Thermo Fisher Scientific, USA) and mounted with Vectashield Antifade mounting medium (Vector Laboratories, USA). Slides were imaged with the EVOS M700 (Thermo Fisher Scientific, USA) at 20x magnification. ImageJ analysis software (National Institutes of Health, Bethesda, USA) was used to quantify the microvessel area within the cortex, CA1, DG, amygdala, thalamus, and striatum. Microvascular coverage was represented as percent of total area (.25 mm^2^).

### Fresh specimen collection and preparation

After neurobehavioral assessment was completed, rats were deeply anesthetized with isoflurane and the brain was collected via decapitation. The hippocampus was isolated, flash frozen in liquid nitrogen, powdered, and stored at −80°C until further usage.

### Western Blot

Powdered hippocampus tissue samples were homogenized in RIPA lysis buffer (Thermo-Scientific, USA) containing protease and phosphatase inhibitor cocktail (Thermo-Scientific, USA). Following lysis using a bead mill homogenizer, the homogenate was centrifuged at 16000 G for 5 minutes at 4°C, supernatants were collected, and total protein concentration was quantified using the BCA protein assay kit (Thermo-Fisher, USA). Protein samples were separated on a 4-20% SDSLJpolyacrylamide gel by electrophoresis (Bio-Rad, USA), according to molecular weight. Proteins were then electroLJtransferred onto polyvinylidene difluoride membranes using the semi-dry transfer method, blocked with 5% skimmed milk at room temperature for 1 hour, and then incubated with primary antibodies ((anti-calcitonin-gene related peptide (CGRP) ms, 1:100, Santa Cruz BioTechnology, USA); (anti-pituitary adenylate cyclase activating peptide (PACAP) rb, 1:5000, Abcam, USA); (anti-endothelial nitric oxide synthase (eNOS) rb, 1:500, Thermo-Fisher, USA); (anti-vascular endothelial growth factor-A (VEGF-A), rb, 1:1000, Proteintech, USA)) overnight at 4°C. After three washes in TBST, the membranes were incubated with secondary (Goat anti-Rabbit-HRP, 1:1000 (Abcam, USA); Goat anti-Mouse HRP, 1:500 (Abcam, USA)) antibodies at room temperature for 1 hour. Signals were detected by chemiluminescence using ECL substrate (Thermo Fisher Scientific, USA) on a BioRad ChemiDoc Imaging System. ImageJ was then used to quantify relative protein levels in the blots. β-actin (Sigma, USA) was used as a loading control for calculation purposes. Western blot data is presented as fold change respective to sham.

### RT-PCR

Total RNA from powdered brain tissue was isolated using Trizol reagent (Life Technologies, Carlsbad, CA). High-capacity cDNA Reverse Transcription Kit (Applied Biosystems, Foster City, CA) was used to synthesize cDNA from the isolated RNA. RT-PCR was performed using the primer sequences as follows: (1) CGRP (F: AGA TGA AAG CCA GGG AGC TG, R: AGG TCT TGT GTG TAC GTG CC), (2) PACAP (F: CAT GTG TAG CGG AGC AAG GTT, R: GTC TTG CAG CGG GTT TCC), (3) eNOS (F: TGA GCA GCA CAA GAG TTA CAA AAT C, R: GCC GCC AAG AGG ATA CCA), (4) VEGF-A (F: GCA CCC ATG GCA GAA GG, R: GGG GTA CCC CTA ACC GCC TCG GCT TGT C), (5) Angiotensin 1 (Ang1) (F: CAC GAC AGA CCA GTA CAA CAC AAA CG, R: GAC GAC TGT TGT TGG TGG TAG CTC T), (6) Tie2 receptor (F: CAG GAC CTT CAC AAC AGC TTC TAT CGG ACT, R: CTG TCG AAG AAT GTC ACT AAG GGT CCA). All primers were obtained from Eurofins Genomics (Louisville, KY). qPCR was performed on a 7500 Real-time PCR System (Applied Biosysems, Foster City, CA) utilizing SYBR Green PCR Master Mix reagents (Applied Biosystems, Foster City, CA). Rat GAPDH (F:AGG TTG TCT CCT GTG ACT TC, R:CTG TTG CTG TAG CCA TAT TC) was used as an endogenous control. The delta-delta calculation method was utilized to obtain fold change relative to controls.

### Statistical Analysis

All data are presented as mean ± standard deviation (SD) and represented visually as bar graphs with individual data points overlaid on top. All statistical analyses were performed using GraphPad Prism (version 9, GraphPad Software, USA). The normal distribution of each variable was verified using both the Shapiro–Wilk test and the Kolmogorov–Smirnov tests. Significant differences were assessed using one-way analysis of variance (ANOVA) followed by Tukey post-hoc test. Data were considered significantly different with α<5% (p<0.05). The correlation coefficient between NOR and Y-maze, and CA1 microvascular constriction, CA1 microvascular degeneration, and MCA wall thickness for sham, 2 weeks 2VO, 4 weeks 2VO, and 6 weeks 2VO were calculated and represented as scatter-plots with best-fit lines superimposed.

## Results

### CCH triggers a gradual decline in cellular density within critical brain regions involved in cognitive processes

Cerebral perfusion was assessed using Laser Speckle Flowmetry before and three days post-2VO surgery. Following 2VO, CBF exhibited a significant decrease of approximately 50% at the three-day mark (**Fig. 1B)**. The density of regional healthy cells experienced a step-wise decrease within the cortex (sham: 322 ± 33; 2wks: 303 ± 31; 4wks: 281 ± 25; and 6wks: 262 ± 36), CA1 (sham: 264 ± 33; 2wks: 189 ± 32; 4wks: 161 ± 47; 6wks: 136 ± 29), amygdala (sham: 275 ± 44; 2wks: 233 ± 20; 4wks: 186 ± 49; 6wks: 179 ± 28), thalamus (sham: 192 ± 32; 2wks: 133 ± 32; 4wks: 129 ± 45; 6wks: 100 ± 9), and striatum (sham: 264 ± 46; 2wks: 167 ± 25; 4wks: 139 ± 51; 6wks: 66 ± 10) (**Fig. 2**). Representative images illustrating cellular health are shown in **SF1**.

**Figure 2.**
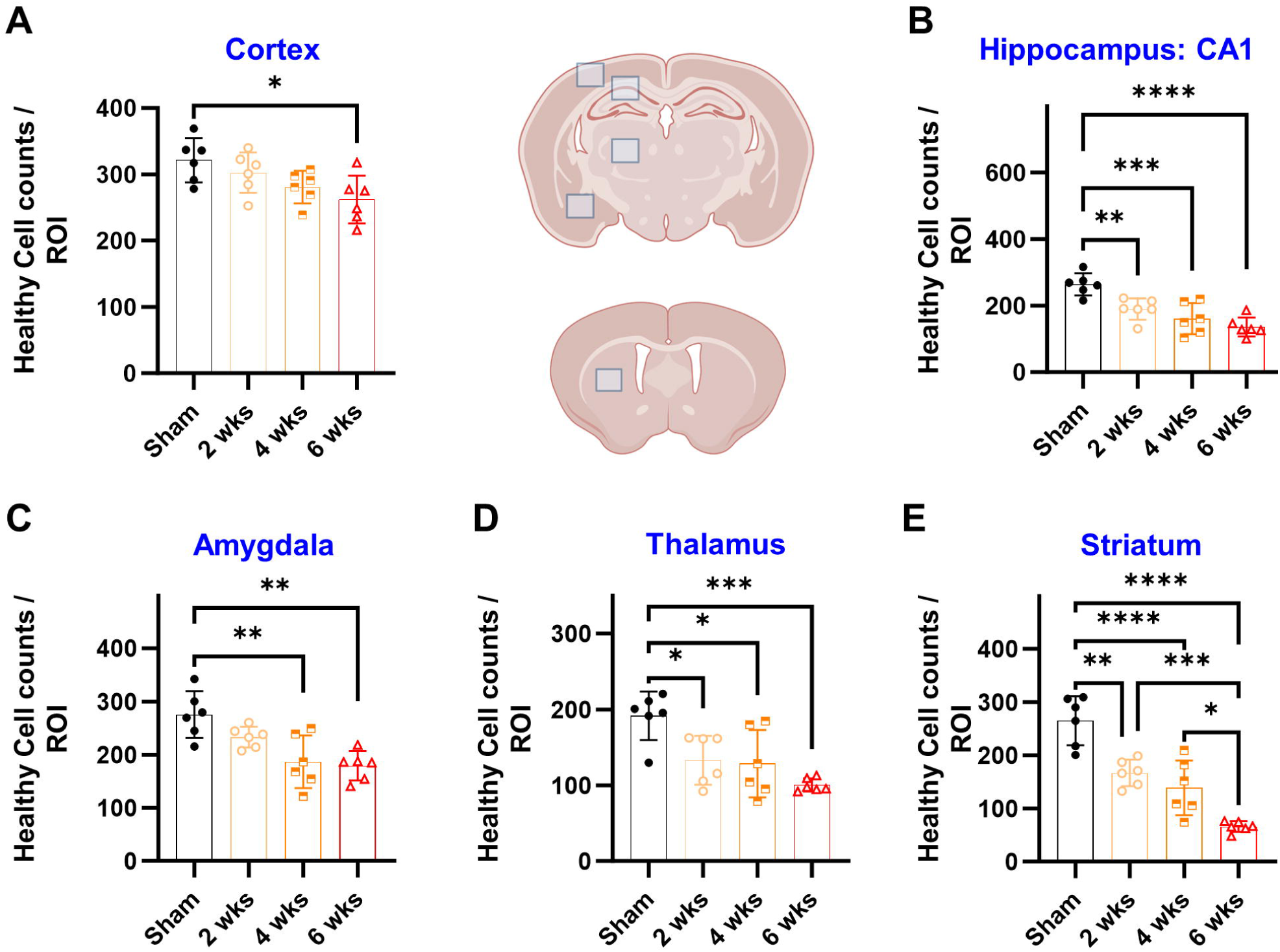
CCH triggers a progressive decline in healthy cell density within brain regions critical to cognitive processes. Healthy cell quantity in a standard ROI (0.25 mm^2^) was measured in the cortex, CA1, amygdala, thalamus, and striatum using semi-automated quantification in ImageJ. Compared to sham, the number of healthy cells in the **(A)** cortex, **(B)** CA1, **(C)** amygdala, **(D)** thalamus, and **(E)** striatum of 2VO rats decreased in a time-dependent manner. (Abbreviations: * *P* < 0.05, ** *P* < 0.01, *** *P* < 0.001, **** *P* <0.0001; 2VO: bilateral common carotid artery occlusion; CCH: chronic cerebral hypoperfusion)

### CCH initiates progressive microvascular constriction and degeneration, while macrovascular function remains relatively unaffected

Macrovascular and microvascular damage were evaluated subsequent to CCH. Initial alterations in MCA and ICA wall thickness returned to baseline levels by the six-week mark post-2VO (**Fig. 3A, SF2**). In contrast, microvascular injury, including both pial and parenchymal constriction, exhibited a significant progressive increase following CCH. Morphological evaluation of cortical pial arterioles post-2VO revealed a rise in the T:D ratio over time (**Fig. 3B**), indicative of time-dependent pial vessel constriction (p<.05). Likewise, gradual increments in parenchymal vessel collapse were noted in the CA1, DG, cortex, thalamus, striatum, and amygdala (**Fig. 3C, SF3**) (p<.05). Further, following CCH, there was evident progression in microvascular degeneration, as denoted by FITC expression. Microvascular coverage decreased significantly in a stepwise manner within the cortex, CA1, striatum, DG, thalamus, and amygdala (**Fig. 4, SF4**) (p<.05).

**Figure 3.**
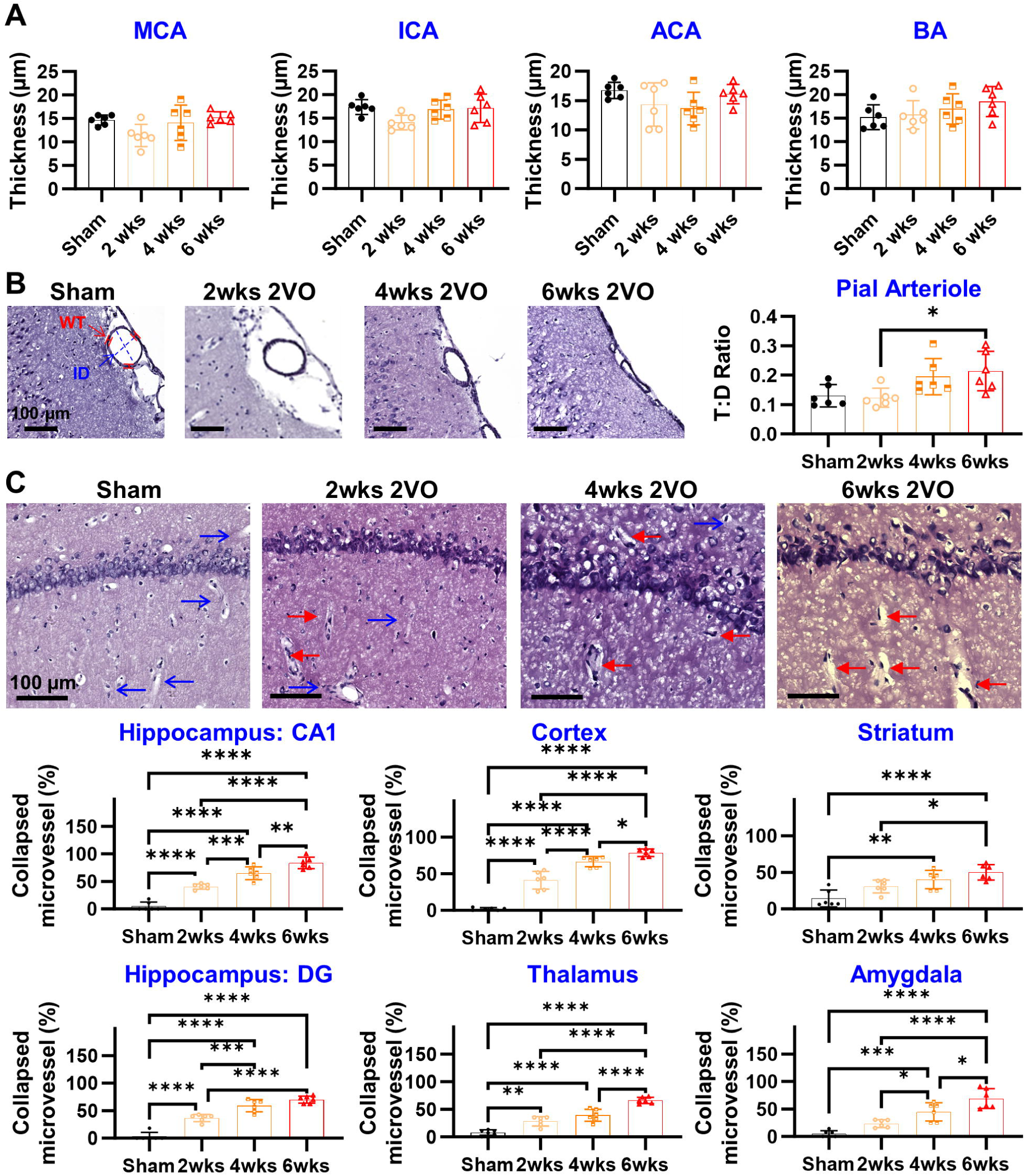
CCH instigates a progressive deterioration in microvascular, but not macrovascular, function. (**A**) Large vessels wall thickness (µm) was measured in the MCA, ICA, ACA, and BA. Early-phase (2wks) morphological changes resolved by late phase (6wks) 2VO. **(B)** The thickness:diameter (T:D) ratio in the cortical pial arterioles increases in a stepwise following 2VO induction, indicating pial vessel constriction (WT, red markers = wall thickness; ID, blue dotted lines and arrows = inner diameter). **(C)** Parenchymal arteriole collapse increased in a time-dependent manner after 2VO in the CA1, cortex, striatum, DG, thalamus, and amygdala. Representative images are from the CA1 of the hippocampus (Blue arrows = healthy vessels; red arrows = constricted vessels). (Abbreviations: * *P* < 0.05, ** *P* < 0.01, *** *P* < 0.001, **** *P* <0.0001; 2VO: bilateral common carotid artery occlusion; ACA: anterior cerebral artery; BA: basilar artery; CCH: chronic cerebral hypoperfusion; ICA: internal carotid artery; MCA: middle cerebral artery)

**Figure 4.**
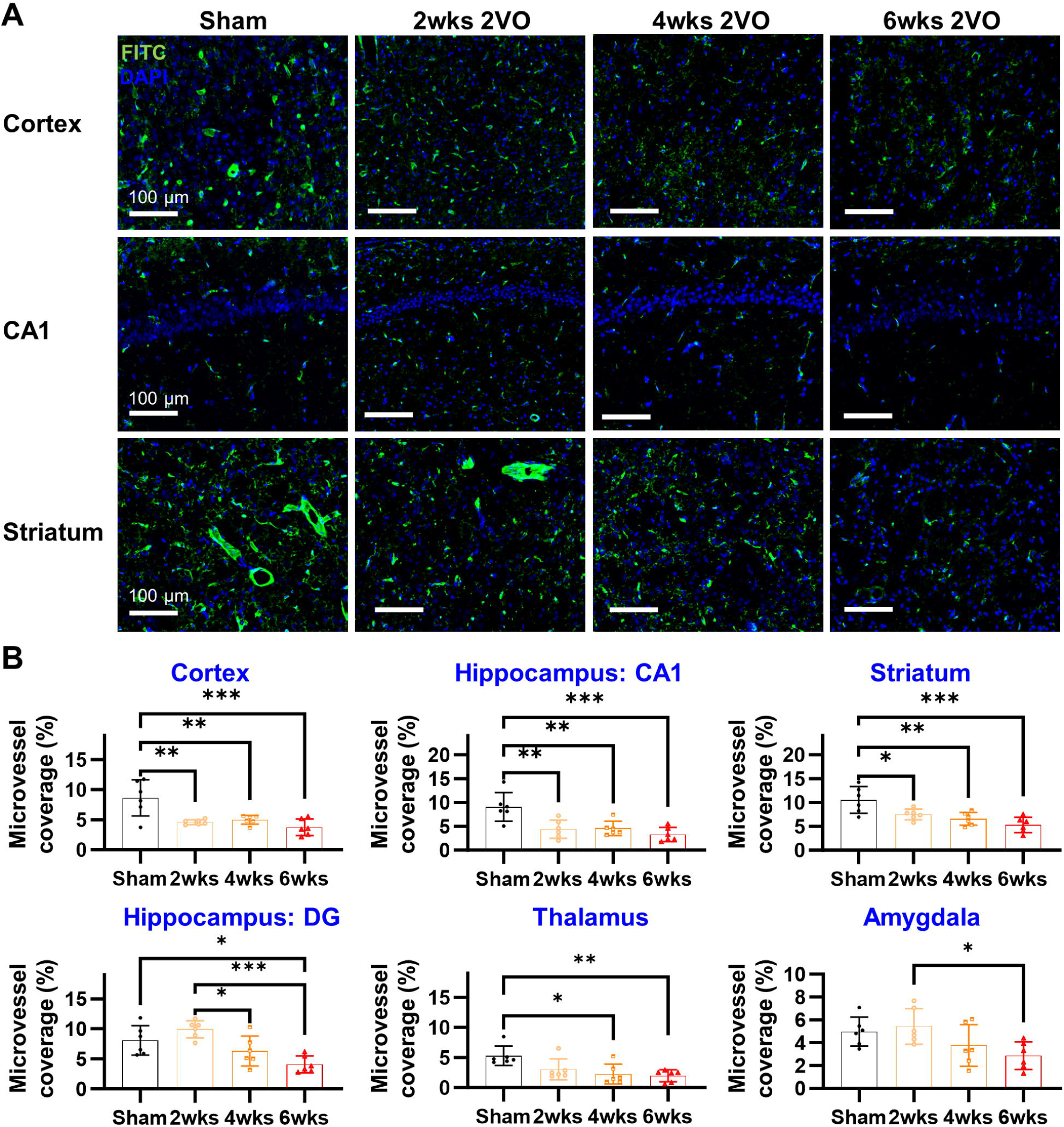
CCH induces progressive microvascular degeneration. (**A**) Microvasculature was imaged using FITC in the cortex, CA1, and striatum. (**B**) Microvascular coverage progressively decreased throughout the brain sub-regions, including the cortex, CA1, striatum, DG, thalamus, and amygdala. (Abbreviations: * *P* < 0.05, ** *P* < 0.01, *** *P* < 0.001; CCH: chronic cerebral hypoperfusion)

### The gradual decline in cognitive function observed in CCH is associated with continuous microvascular damage

Both working recognition memory and long-term spatial memory were assessed in connection with the documented microvascular dysfunction. The 2VO model resulted in a gradual deterioration of working recognition memory (sham: 0.34 ± 0.22; 2wks: −0.01 ± 0.24; 4wks: −0.14 ± 0.22; and 6wks: −0.35 ± 0.27) (**Fig. 5A**) and long-term spatial memory (sham: 0.46 ± 0.14; 2wks: 0.36 ± 0.08; 4wks: 0.34 ± 0.08; and 6wks: 0.27 ± 0.11) (**Fig. 5B**).

**Figure 5.**
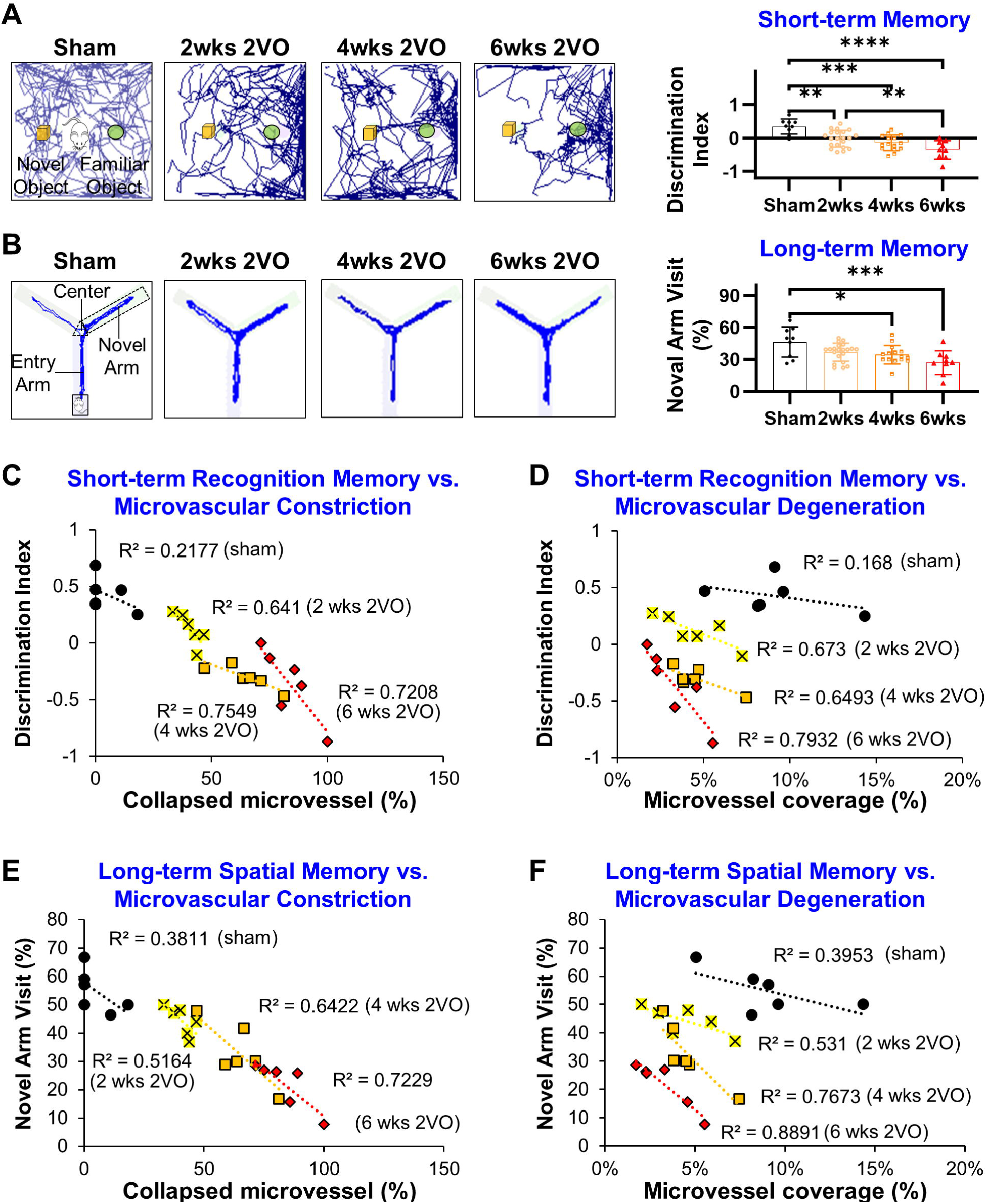
Gradual decline in cognitive function correlates with continuous microvascular damage in CCH. **(A)** Novel object recognition discrimination index decreased in a stepwise manner, indicating loss of working memory. (**B)** Long-term spatial memory decreased in a progressive manner, as assessed by the amount of novel arm visits in the Y-maze. (**C**) Short-term recognition memory has the strongest correlation with microvascular constriction, (**D**) followed by microvascular degeneration. (**E & F**) Long-term spatial memory also correlated strongly with microvascular constriction and degeneration. (Abbreviations: * *P* < 0.05, ** *P* < 0.01, *** *P* < 0.001, **** *P* <0.0001; CCH: chronic cerebral hypoperfusion)

The correlation between NOR results and MCA wall thickness, CA1 vascular constriction, and CA1 vascular degeneration was computed. The NOR was selected as the primary assessment of cognitive dysfunction due to the relevance to short-term memory in VCI [46]. Consistent with our previous findings indicating microvascular, rather than macrovascular, impairment as the primary determinant, macrovascular wall thickness exhibited the lowest correlation coefficient (**SF5**). Microvascular constriction demonstrated the highest correlation coefficient (sham: R^2^=0.2177, 2wks: R^2^=0.641, R^2^=0.7549, R^2^=0.7208) (**Fig. 5C**) closely followed by microvascular degeneration (sham: R^2^=0.168, 2wks: R^2^=0.673, R^2^=0.6493, R^2^=0.7932) (**Fig. 5D**). Y-maze correlation was also evaluated as a secondary measure of cognitive impairment [47], and although it also displayed a strong correlation to microvascular dysfunction (**Fig. 5E & F**), the NOR consistently exhibited the most robust correlations (**Fig 5C**).

### DR triggers the upregulation of vasodilatory mediators

To gauge DR’s vasodilatory potential in CCH, we investigated the hippocampal expression of vasodilatory mediators including CGRP, PACAP, and eNOS. We observed a decrease in hippocampal CGRP expression at 6 wks post-2VO induction (sham: 1±0.29; 6wks 2VO: 0.81±0.76), which was significantly upregulated by DR treatment (6wks 2VO+DR: 1.68±0.73) (**Fig. 6A**). Similarly, PACAP expression in the hippocampus markedly decreased at 6wks post-2VO induction, and this trend was significantly reversed by DR treatment (**Fig. 6B**). Additionally, we noted a significant decrease in hippocampal eNOS expression at 6wks post-2VO induction (sham: 1.00±0.12; 6wks 2VO: 0.21±0.16), which was prevented by DR treatment (6wks 2VO+DR: 1.17±0.38) (**Fig. 6C**). mRNA levels of CGRP, PACAP, and eNOS exhibited similar trends to protein expression (**Fig. 6D, 6E, 6F**). No significant differences were noted in the 2VO animals treated with swimming compared to those subjected to 2VO alone.

**Figure 6.**
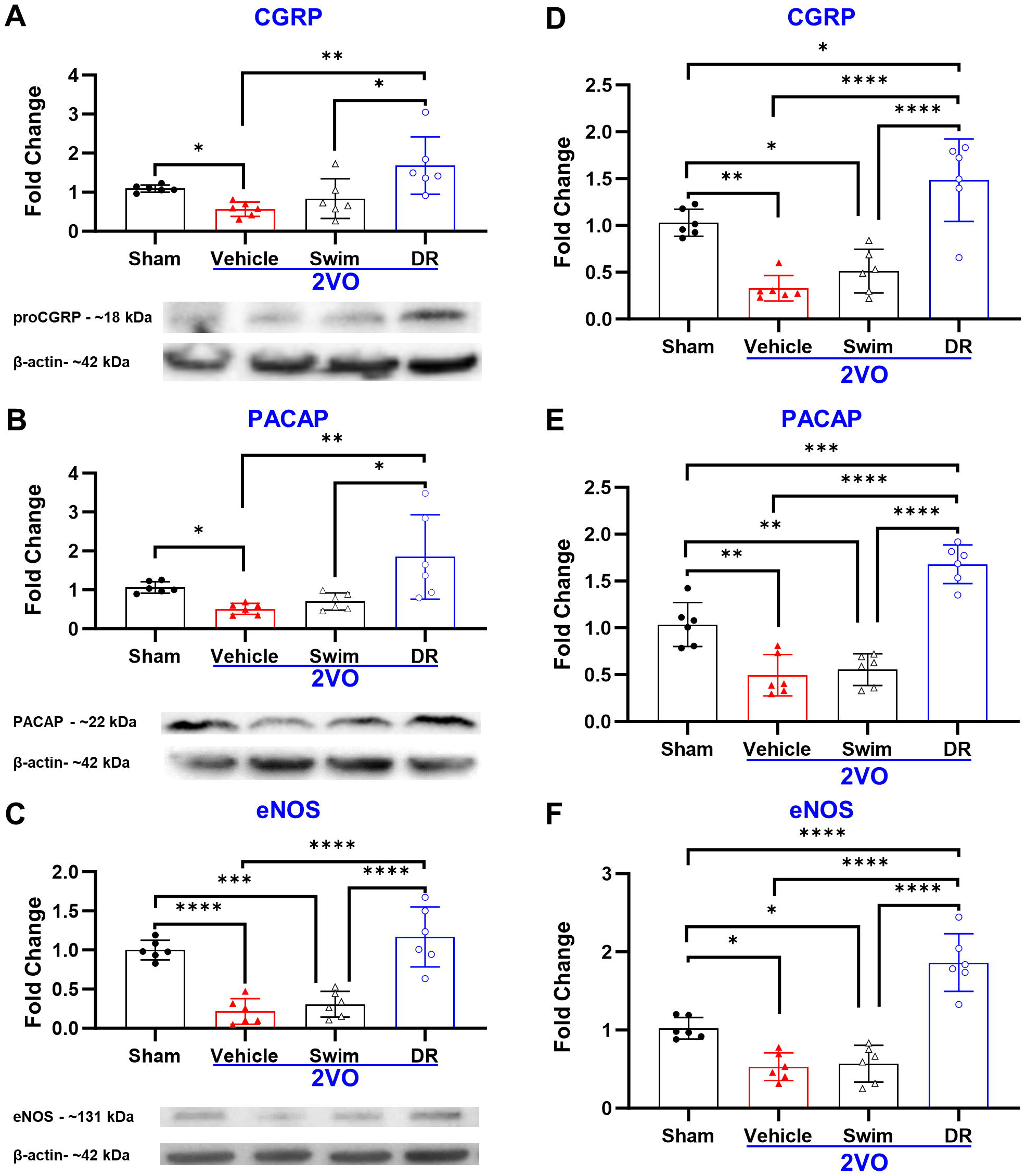
DR stimulates vasodilatory factors in CCH at 6wks. Protein expression and mRNA levels of vasodilatory factors in the hippocampus were assessed at 6wks. DR prevented 6wk downregulation of (**A & D**) CGRP, (**B & E**) PACAP, and (**C & F**) eNOS. (Abbreviations: * *P* < 0.05, ** *P* < 0.01, *** *P* < 0.001, **** *P* <0.0001; 2VO: bilateral common carotid artery occlusion; CCH: chronic cerebral hypoperfusion; CGRP: calcitonin gene related peptide; DR: diving reflex; eNOS: endothelial nitric oxide synthase; PACAP: pituitary adenylate cyclase activating peptide)

### DR triggers the upregulation of angiogenic mediators

We evaluated hippocampal expression of VEGF-A to gauge DR’s angiogenic potential. VEGF-A expression was significantly reduced in the 2VO group (sham: 1.00±0.12; 6wks 2VO: 0.59±0.17) (**Fig. 7A**). However, DR application led to an upregulation of VEGF-A expression (6wks 2VO+DR: 1.58±0.55) compared to both the 2VO and sham groups. mRNA levels of VEGF-A, angiopoietin-1, and Tie2 mirrored the trends observed in VEGF-A protein expression, with notable downregulation in the 2VO group that was ameliorated by DR (**Fig. 7B, 7C, 7D**). There were no significant differences observed between the 2VO animals treated with swimming and those subjected to 2VO alone.

**Figure 7.**
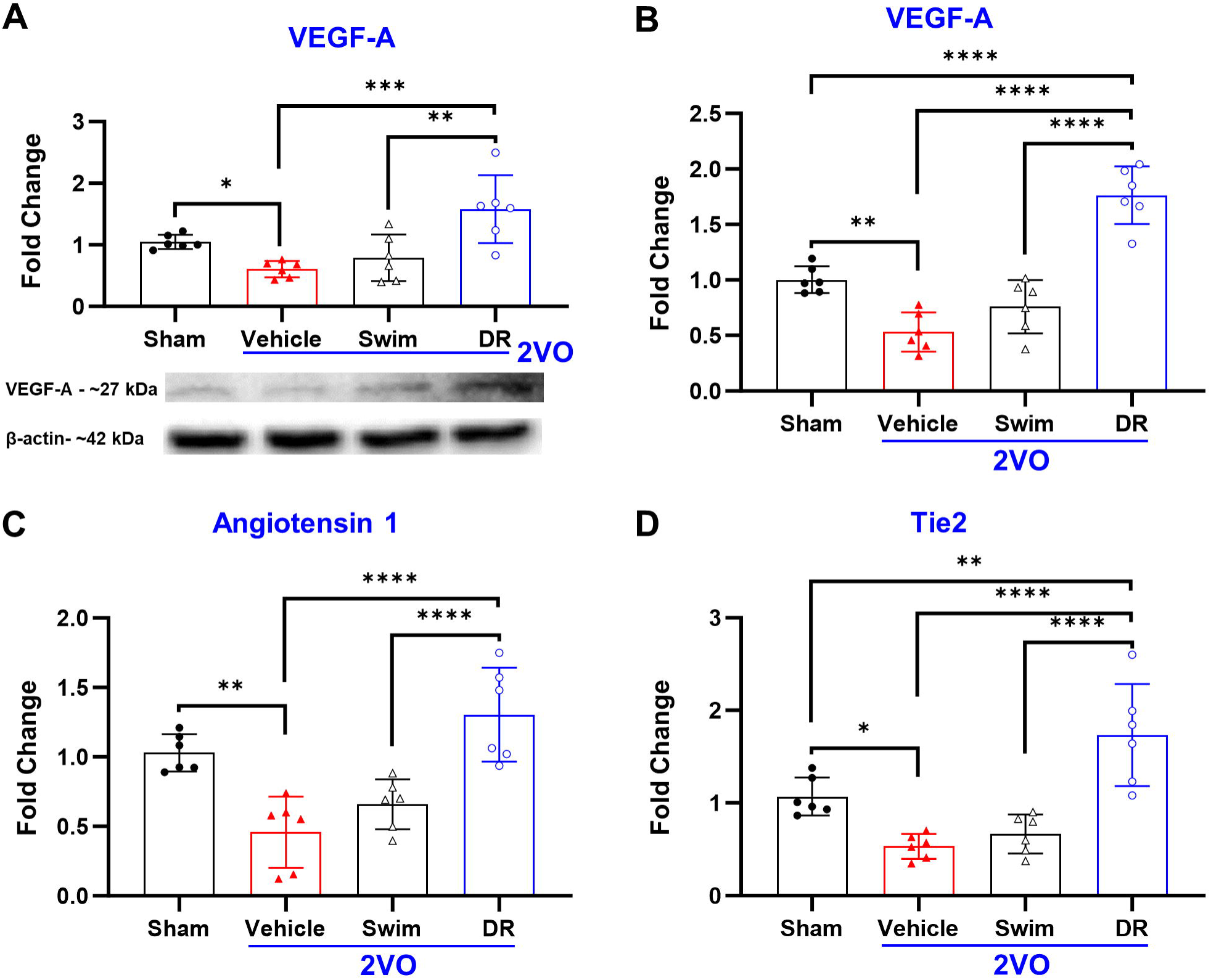
DR stimulates angiogenic factors in CCH at 6wks. Protein expression and mRNA levels of angiogenic factor in the hippocampus was assessed at 6wks. DR promoted (**A & B**) VEGF-A, (**B**) Angiotension, and (**C**) Tie2 expression. (Abbreviations: * *P* < 0.05, ** *P* < 0.01, *** *P* < 0.001; 2VO: bilateral common carotid artery occlusion; CCH: chronic cerebral hypoperfusion; DR: diving reflex; VEGF-A: vascular endothelial growth factor-A)

### DR enhances short-term recognition memory by ameliorating microvascular dysfunction in the cortex and hippocampal subfields

DR decreased the pial thickness-to-diameter (T:D) ratio, restoring it to levels comparable to sham-operated animals (sham: 0.12±0.03 µm, 6wks 2VO: 0.22±0.07 µm vs. 6wks 2VO+DR: 0.11±0.01 µm) (**Fig. 8A**). Similarly, DR prevented constriction of parenchymal microvessels in the cortex (sham: 3.38%±2.19%, 6wks 2VO: 83.92%±6.04% vs. 6wks 2VO+DR: 47.52%±8.66%), CA1 (sham: 3.36%±5.25%, 6wks 2VO: 95.83%±6.45% vs. 6wks 2VO+DR: 33.80%±9.11%), and DG (sham: 0%, 6wks 2VO: 88.13%±14.73% vs. 6wks 2VO+DR: 51.68%±5.22%) (**Fig. 8B**). Additionally, DR alleviated microvascular degeneration within the cortex (sham: 8.1%±1.8%; 6wks 2VO: 3.0%±0.9% vs. 6wks 2VO+DR: 8.9%±3.1%), CA1 (sham: 8.0%±1.7%; 6wks 2VO: 2.8%±1.3% vs. 6wks 2VO+DR: 10.1%±2.4%), and DG (sham: 8.5%±2.4%; 6wks 2VO: 4.5%±2.3% vs. 6wks 2VO+DR: 11.6%±2.1%) (**Fig. 8C**). The improvement in microvascular function induced by DR was accompanied by significantly enhanced working recognition memory (sham: 0.33±0.26; 6wks 2VO: −0.39±0.26 vs. 6wks 2VO+DR: 0.12±0.11) (**Fig. 8D**). Swimming treatment did not provide any improvement over 2VO alone in either cortical and hippocampal microvascular dysfunction or short-term memory.

**Figure 8.**
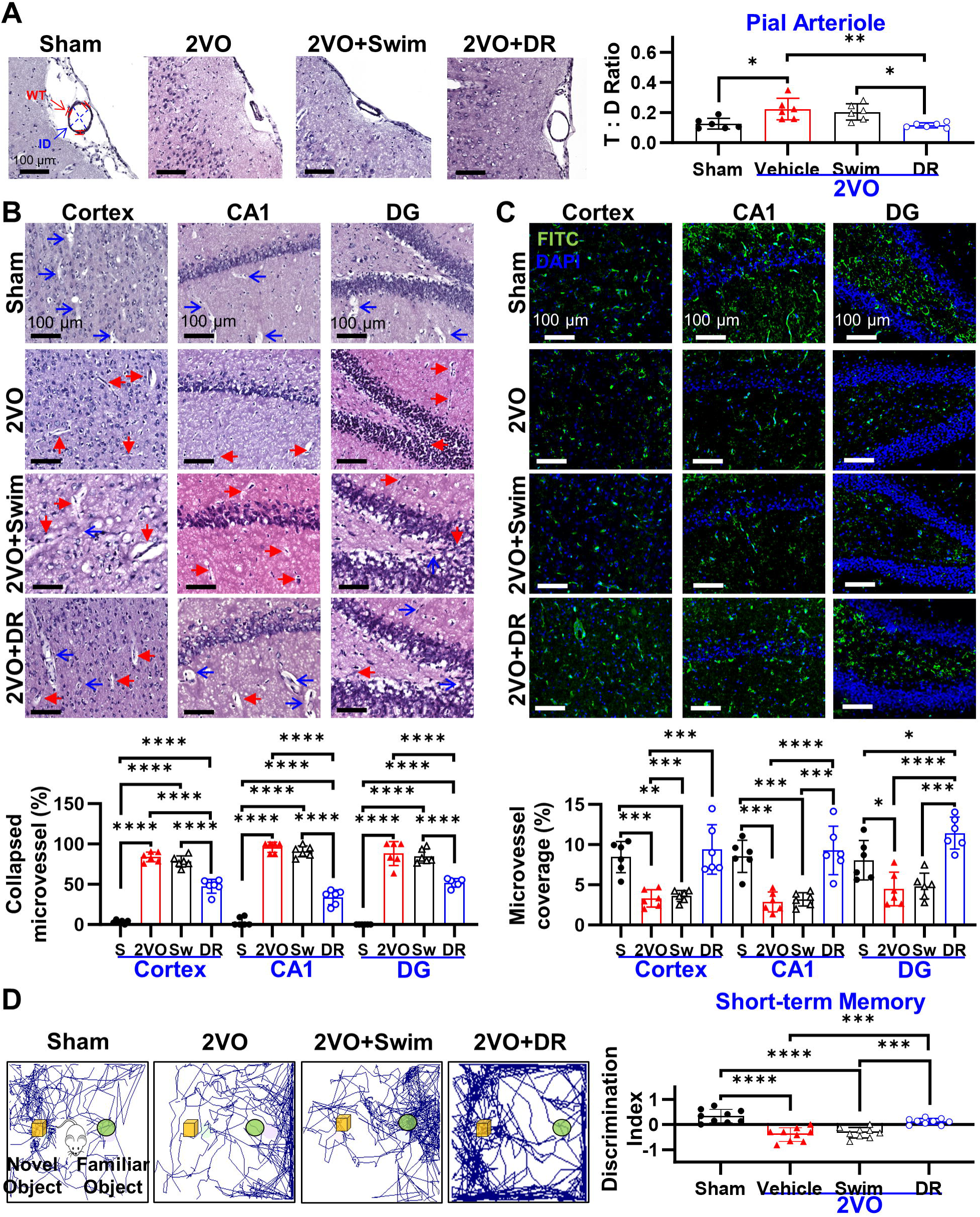
DR boosts short-term recognition memory by ameliorating cortical and hippocampal microvascular dysfunction at 6wks. **(A)** The thickness:diameter (T:D) ratio in the cortical pial arterioles decreased with the application of chronic DR, indicating a decrease of pial constriction (WT, red markers = wall thickness; ID, blue dotted lines and arrows = inner diameter). **(B&C)** Cortical and hippocampal parenchymal arteriole collapse and microvascular degeneration were prevented with DR (Blue arrows = healthy vessels; red arrows = constricted vessels). **(D)** Novel object recognition discrimination index increased with DR treatment, indicating improved working memory. (Abbreviations: * *P* < 0.05, ** *P* < 0.01, *** *P* < 0.001, ****( *P* < 0.0001; CCH: chronic cerebral hypoperfusion; DR: diving reflex)

### DR facilitates long-term spatial memory by mitigating microvascular dysfunction in the amygdala and white matter

DR also alleviated microvascular constriction and microvascular degeneration in the amygdala, thalamus, and striatum. DR significantly prevented parenchymal vessel collapse in the amygdala (sham: 7.07%±6.19%, 6wks 2VO: 68.35%±19.54% vs. 6wks 2VO+DR: 35.81%±7.81%), thalamus (sham: 9.36%±3.78%, 6wks 2VO: 71.33%±8.66% vs. 6wks 2VO+DR: 39.11%±12.17%), and striatum (sham: 8.12%± .07%, 6wks 2VO: 43.28%±7.61% vs. 6wks 2VO+DR: 27.74%±7.94%) (**Fig. 9A**). Additionally, DR ameliorated microvascular degeneration in the amygdala (sham: 4.5%±0.8%; 6wks 2VO: 2.8%±1.3% vs. 6wks 2VO+DR: 7.5%±3.8%), thalamus (sham: 4.6%±0.3%; 6wks 2VO: 2.3%±1.4% vs. 6wks 2VO+DR: 6.1%±1.3%), and striatum (sham: 10.8%±2.0%; 6wks 2VO: 5.7%±1.4% vs. 6wks 2VO+DR: 9.0%±1.6%) (**Fig. 9B**). Swimming did not prevent either parenchymal constriction or microvascular degeneration within the amygdala, thalamus, and striatum. The microvascular enhancement induced by DR was associated with significant improvements in long-term spatial memory (sham: 0.48±0.19; 6wks: 0.24±0.10, vs. 6wks 2VO+DR: 0.37±0.09) (**Fig. 9C**); swimming provided no discernible benefit to long-term spatial memory.

**Figure 9.**
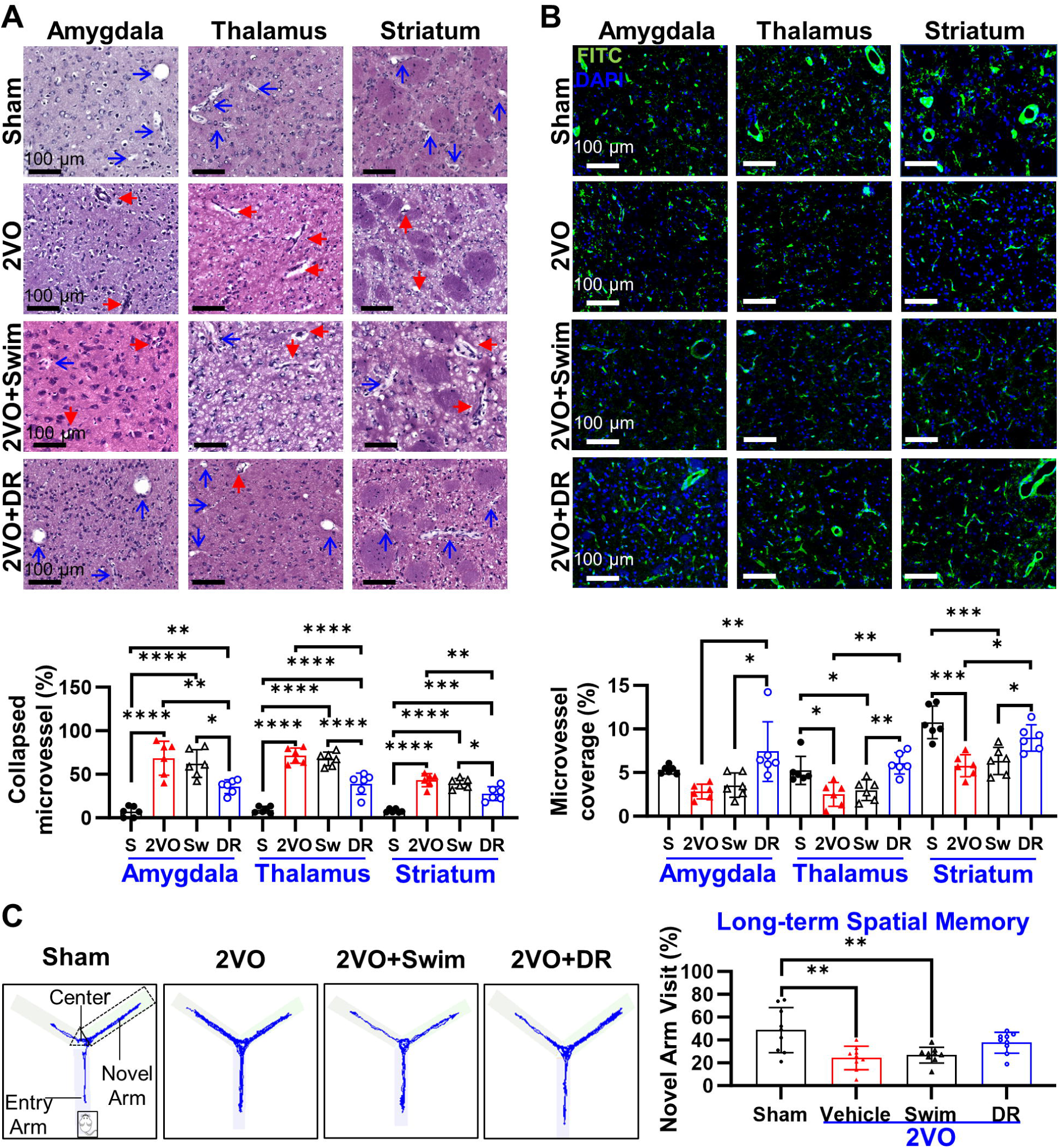
DR promotes long-term spatial memory by diminishing microvascular injury in the amygdala and white matter at 6wks. **(A&B)** Parenchymal collapse and microvascular loss was prevented in the amygdala, thalamus, and striatum with DR treatment **(C)** Long-term spatial memory function increased with DR, as assessed by the amount of novel arm visits in the Y-maze. (Blue arrows = healthy vessels; red arrows = constricted vessels). (. Abbreviations: * *P* < 0.05, ** *P* < 0.01, *** *P* < 0.001, **** *P* < 0.0001; CCH: chronic cerebral hypoperfusion; DR: diving reflex)

## Discussion

To the best of our knowledge, this study represents one of the few instances of evidence suggesting a notable correlation between progressive memory dysfunction and the incremental constriction and degeneration of cerebral microvasculature following CCH. These findings underscore the pivotal role of microvascular injury in the initiation and advancement of cognitive decline in the context of CCH. Therefore, therapeutic strategies aimed at mitigating microvascular injury may hold promise as effective disease-modifying approaches in VCI. Our experiments demonstrate that DR provides significant protection to the microvasculature by reducing microvascular constriction and promoting extensive angiogenesis, particularly observed in the hippocampus and white matter (**Fig. 10**). This preservation leads to notable retention of short-term working recognition and long-term spatial memory capacities. Based on our current understanding, this study represents the first documentation of DR’s preclinical efficacy in VCI, thereby furnishing evidence for DR as a promising non-pharmacological intervention capable of directly safeguarding the microvasculature and mitigating cognitive impairment induced by CCH.

**Figure 10.**
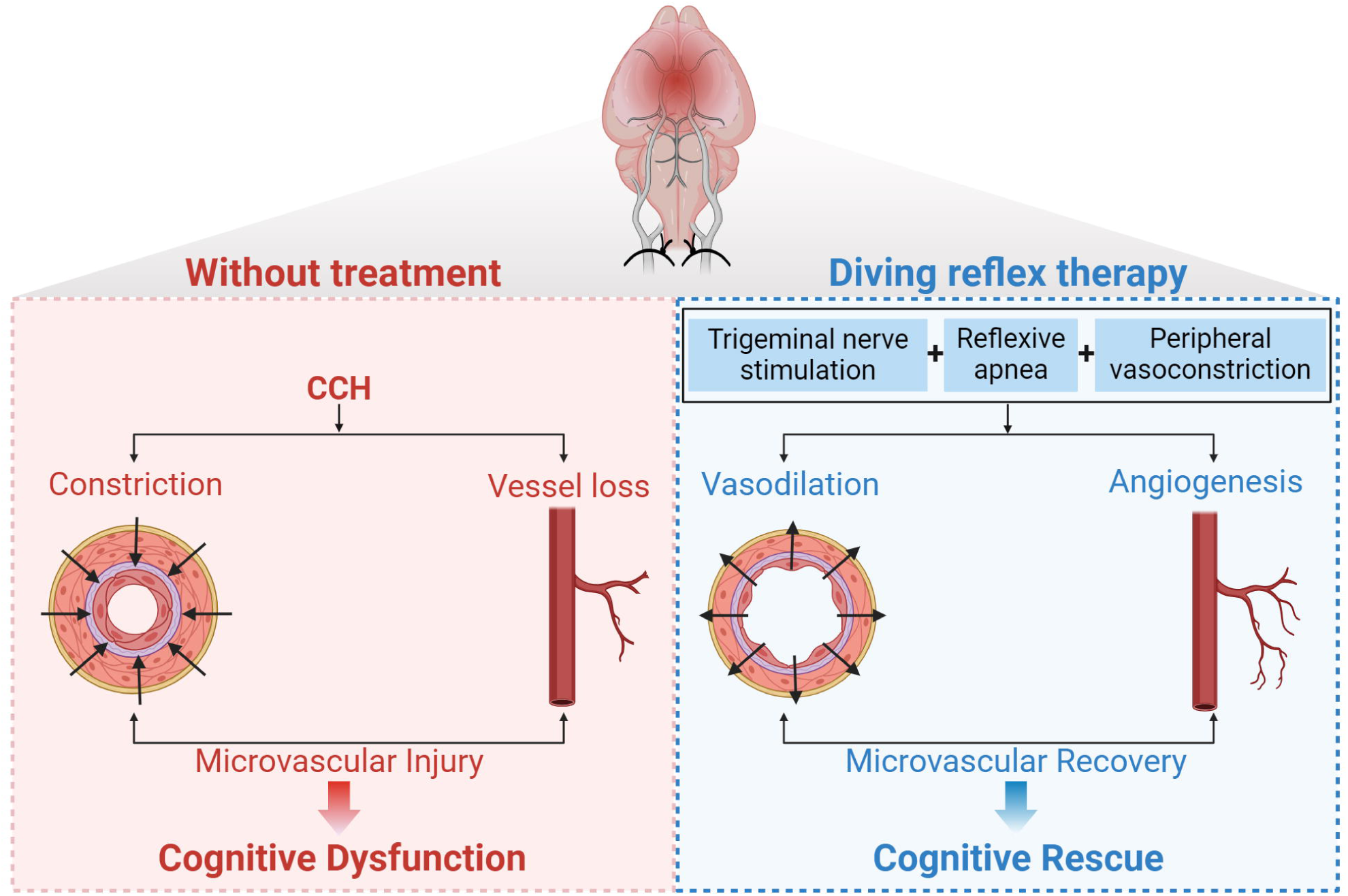
Intrinsic DR enhances cognitive performance by mitigating microvascular dysfunction in vascular cognitive impairment. Vascular cognitive impairment is due to CCH- induced vasoconstriction and vessel loss, which together lead to microvascular injury. Via the combined mechanisms of vasodilation and angiogenesis, DR encourages microvascular recovery. This promotes cognitive improvement and prevents vascular cognitive impairment. (Abbreviations: CCH: chronic cerebral hypoperfusion; DR: diving reflex)

Our investigation revealed that cognitive dysfunction induced by CCH primarily stems from microvascular dysfunction rather than large vessel injury. At the late stage of CCH, six weeks post-2VO, any morphological alterations observed in large vessels at two weeks had resolved, consistent with prior findings. Although decreases in large vessel wall thickness at two weeks might serve as a protective mechanism [48], evaluations at subsequent time intervals verify the lack of macrovascular changes by six weeks [49–51]. Conversely, CCH induces a progressive microvascular injury characterized by microvascular constriction and degeneration, strongly associated with progressive memory decline. 2VO initiates a continual constriction of microvessels and a decrease in microvascular coverage, resulting in a significant 60% reduction in vasculature within the CA1 region at six weeks. Importantly, microvascular damage, as opposed to macrovascular damage, correlates with gradual impairments in both short-term recognition and long-term spatial memories, leading to a maximal loss of 50% in memory capacity. Our findings are consistent with prior research demonstrating microvascular thinning and shortening [15], decreased perfused capillaries and microvascular density [11,49–52], and diminished vasodilatory response [14] at the later stage of 2VO. This study underscores the importance of microvascular dysfunction in VCI, elucidating the relationship between its gradual progression and the development of cognitive dysfunction.

DR is identified as a distinctive intrinsic mechanism with the capacity to trigger a robust cerebral vasodilatory response [20,28], thereby offering a potential therapeutic pathway for CCH. This investigation unveils, for the first time, that DR conserves microcirculation by obstructing microvascular constriction, accomplished through a significant augmentation in vasoactive neuropeptides, including CGRP and PACAP, compared to 2VO alone, culminating in a substantial 0.64-fold reduction in vascular constriction within the CA1 region. Activation of the trigeminal nerve induces the release of multiple vasoactive neuropeptides [32,55], which have been shown to exhibit vasodilatory effects at the microvessel level [54–57]. Moreover, DR notably amplifies expression of eNOS, which is indispensable for preserving vasodilatory capacity [58], compared to other pharmacological interventions like angiotensin-(1-7) [59], by approximately 5.5-fold, relative to the state of 2VO alone. Notably, DR exerts a substantial effect even when contrasted with non-pharmacological interventions like physical exercise and RLIC, which augment eNOS expression by approximately 3.5-fold in the hippocampus [30,60]. To the best of our knowledge, this investigation represents the initial attempt to specifically address microvascular constriction with the aim of augmenting cognitive function in VCI. This pioneering therapeutic approach underscores the potential of DR as an exceedingly promising intervention strategy.

In addition to its vasodilatory effects, DR further facilitated notable angiogenesis, resulting in an approximately 3-fold increase in vascular coverage within the CA1 region. This coincided with a significant upregulation of angiogenesis-related genes such as VEGF-A, angiopoietin-1, and Tie2. Comparative analysis with other non-pharmacological interventions suggests that DR may demonstrate an augmented capacity for preserving vascular coverage. RLIC enhances hippocampal vasculature by approximately 1.3-fold in CCH [30], while early aerobic exercise does not elicit significant alterations in functional capillary density in the cortex [51]. The elevation of VEGF-A levels induced by DR may contribute to the observed enhancement in microvascular coverage; VEGF-A stimulates endothelial cell proliferation and migration, thereby fostering neovascularization [50]. While alternative interventions such as enriched environment [61] and transcranial magnetic stimulation [62] also elevate VEGF-A levels in CCH, DR-mediated VEGF elevation surpasses sham levels, suggesting a potentially more robust angiogenic effect. However, despite RLIC also elevating VEGF levels beyond sham levels [13], it fails to surpass sham levels in terms of vascular coverage [13,30], as observed with DR. This implies that while RLIC and DR may share the mechanism of VEGF elevation mediated by peripheral constriction [19], the effects of DR are not solely dependent on VEGF. Consequently, further exploration of the diverse mechanisms underlying DR’s angiogenic effect holds substantial promise in the context of CCH.

Among the various interventions in CCH, targeting microvascular function has shown significant promise in improving cognitive function. DR, which concurrently addresses multiple mechanisms to preserve microvascular function, leads to notable cognitive enhancements. It enhances working memory by approximately 86% and long-term spatial memory by around 56% compared to 2VO alone. As a vasodilatory agent, nimodipine has been observed to enhance learning and memory by about 60% in 2VO rats, by suppressing autophagy in the CA1 and CA3 regions [63]. Induction of vasodilation in the vertebral arteries by prostaglandin E1 and DI-3-n-Butylphthalide also improves learning and memory by approximately 80% [11,12], linked with VEGF upregulation in CCH [12,50]. Similarly, preservation of vascular density and BBB integrity underscores the approximately 75% improvement in memory with methylene blue treatment [64], while promotion of angiogenesis through platelet-rich plasma and epidermal neural crest stem cell transplantation enhances neurobehavior [65]. Moreover, enhancement of angiogenesis mediated by eNOS activation induced by RLIC improves learning memory by about 60% in CCH [30]. Our findings suggest that DR harnesses all these factors, resulting in a significant improvement in cognition comparable to that induced by other preclinical therapeutics targeting microvascular dysfunction.

From a mechanistic perspective, the effects of DR may be attributed to a combination of trigeminal nerve stimulation, apnea, and peripheral vasoconstriction (**Fig 10**). While physical activities such as running and swimming can induce an angiogenic response, these interventions necessitate a certain level of physical exertion to achieve efficacy [15,51,52,66–69]. In human subjects, only high-intensity physical exercise, not moderate-intensity exercise, promotes an elevation in VEGF and nitric oxide, a key vasodilator [70]. Similarly, in healthy animal models, rigorous exercise administered at regular intervals augments vascular coverage within the cortex [71], whereas intermittent running does not impact the number of string vessels in the corpus callosum [68]. Regarding swimming, only mild enhancements in brain-derived neurotrophic factor, and hence angiogenic capacity, are noted with moderate-intensity training in rats with vascular dementia, compared to the restoration to healthy levels observed with high-intensity swimming [72]. It is noteworthy that the rats subjected to 2VO, either with DR or swim, did not engage in either high-intensity or prolonged physical exercise. Instead, they underwent brief bouts (less than 30 seconds) with a 5-minute interval between each bout. Consequently, the beneficial effects of DR, which were absent in animals treated with swimming, can be attributed to, and underscore the significance of, the synergistic effects stemming from trigeminal nerve stimulation, apnea, and peripheral vasoconstriction.

This study has some limitations. Firstly, only a single regimen of DR was administered in this evaluation. While it demonstrated significant microvascular protection leading to notably improved cognitive function, the exploration of alternative DR dosages may unveil a more profound protective effect in CCH. Therefore, further investigation into the dose-response relationship of DR in CCH is warranted. Secondly, a comprehensive understanding of the intricate mechanistic pathways governing microvascular function remains lacking. Despite the evidence of DR triggering vasodilation and angiogenesis, and modulating associated vasodilatory and angiogenic factors, the precise underlying cellular and molecular mechanisms remain to be elucidated. Thirdly, memory was the sole measure of cognitive dysfunction assessed in this study, reflecting the most commonly used assessment in CCH research. However, CCH impacts various other forms of behavior, suggesting that employing a broader array of behavioral assessments, encompassing anxiety- and depression-like behaviors, may enhance the translation of DR to clinical applications.

The administration of DR three days following 2VO (during the early phase of VCI) effectively mitigates microvascular dysfunction and enhances memory performance, indicating its potential as an innovative therapeutic approach for CCH. Notably, the simultaneous augmentation of microvascular vasodilation and angiogenesis suggests that DR may not only arrest the progression of CCH but also promote microvessel restoration and reverse disease progression. These findings strongly support the use of DR as a disease-modifying treatment for CCH-induced microvascular impairment. In contrast to conventional vasodilators, which may induce systemic hypotension, DR-induced peripheral vasoconstriction could mitigate this potential side effect. This unique advantage positions DR as a promising standalone and/or adjunctive therapeutic strategy for VCI. DR has demonstrated clinical efficacy in assessing autonomic nervous system disorders [73] and managing supraventricular tachycardia through facial immersion in cold water [74–76]. Cold facial immersion has emerged as a promising method for inducing DR and has shown the ability to increase CBF in humans [26,27], suggesting its potential as a non-invasive therapeutic approach for cerebral hypoperfusion disorders such as VCI. Overall, the non-invasive nature, targeted effects, and established safety profile of DR make it an attractive candidate for clinical translation in the treatment of CCH-induced VCI.

## Conclusion

The findings of this study demonstrate, for the first time, the significant protective influence of DR on the microvasculature in CCH, particularly notable within the hippocampus and white matter regions of the brain. The combined enhancement of microvascular vasodilation and angiogenesis suggests that DR may not only impede the progression of CCH but also facilitate microvessel repair and reverse disease advancement. This marks a significant paradigm shift in VCI management, diverging from conventional approaches primarily focused on symptomatic relief. The synergistic cerebrovascular benefits, facilitated through trigeminal nerve stimulation and peripheral vasoconstriction, led to significant improvements in memory function, crucial clinical markers of VCI severity and progression. Furthermore, the promising underlying mechanisms involved in preserving microvasculature integrity hold potential for extending benefits to other conditions characterized by cerebral hypoperfusion, such as Alzheimer’s disease and Parkinson’s disease. Combining the non-pharmacological and non-invasive attributes of DR with its profound protective impacts on microvasculature underscores its potential as an effective therapeutic intervention for a spectrum of diseases impacted by CCH.

## Figure Legends

**Supplementary Figure 1. CCH induces a progressive decline in healthy cell density in key brain areas affecting cognition**. Representative images of ImageJ quantification in sham and 2wks, 4wks, and 6wks 2VO (black bars = 100 µm). (Abbreviations: 2VO: bilateral common carotid artery occlusion; CCH: chronic cerebral hypoperfusion)

**Supplementary Figure 2. CCH does not induce macrovascular dysfunction.** Large vessel wall thickness (µm) was measured in the MCA, ICA, ACA, and BA. Representative images of vessel thickness in sham and 2wks, 4wks, and 6wks 2VO are taken at 20X. (Abbreviations: 2VO: bilateral common carotid artery occlusion; ACA: anterior cerebral artery; BA: basilar artery; CCH: chronic cerebral hypoperfusion; ICA: internal carotid artery; MCA: middle cerebral artery)

**Supplementary Figure 3. CCH induces progressive microvascular constriction**. Parenchymal arteriole constriction in the cortex, striatum, DG, thalamus, and amygdala increased in a time-dependent manner after 2VO (Blue arrows = healthy vessels; red arrows = constricted vessels). (Abbreviations: 2VO: bilateral common carotid artery occlusion; CCH: chronic cerebral hypoperfusion; DG: dentate gyrus)

**Supplementary Figure 4. CCH induces stepwise microvascular degeneration.** Microvascular degeneration occurred in a stepwise manner throughout the DG, thalamus, and amygdala after 2VO. (Abbreviations: 2VO: bilateral common carotid artery occlusion; CCH: chronic cerebral hypoperfusion; DG: dentate gyrus)

**Supplementary Figure 5. Memory impairment in CCH does not exhibit a significant correlation with macrovascular dysfunction**. Short-term recognition memory and long-term spatial memory show no correlation with MCA wall thickness in 2VO. (Abbreviations: 2VO: bilateral common carotid artery occlusion; CCH: chronic cerebral hypoperfusion; MCA: middle cerebral artery)

## Acknowledgments

This work is supported in part by career enhancement award at Feinstein Institutes for Medical Research and the Zoll Foundation Grant. The authors would like to thank Drs. Betty Diamond, Patricio Huerta, Valentin Pavlov, Annette Lee, Christine Metz, and Barbara Sherry for their insightful evaluation and comments on the manuscript.

## Author contributions

C.L. conceptualized and designed the experiments. W.T., K.P., S.W., and C.L. performed experiments. W.T., K.P., S.W., and C.L. analyzed the data. C.L. acquired funding for the project. W.T. and C.L. prepared the manuscript. K.P., S.W., D.S., M.B., and C.L. critically revised the manuscript.

## Data and materials availability

The datasets used and/or analyzed during the current study are available from the corresponding author upon reasonable request.

