## Supplementary figures and images for "Intrinsic diving reflex enhances cognitive performance by alleviating microvascular dysfunction in vascular cognitive impairment"

### Supplemental Figure 1

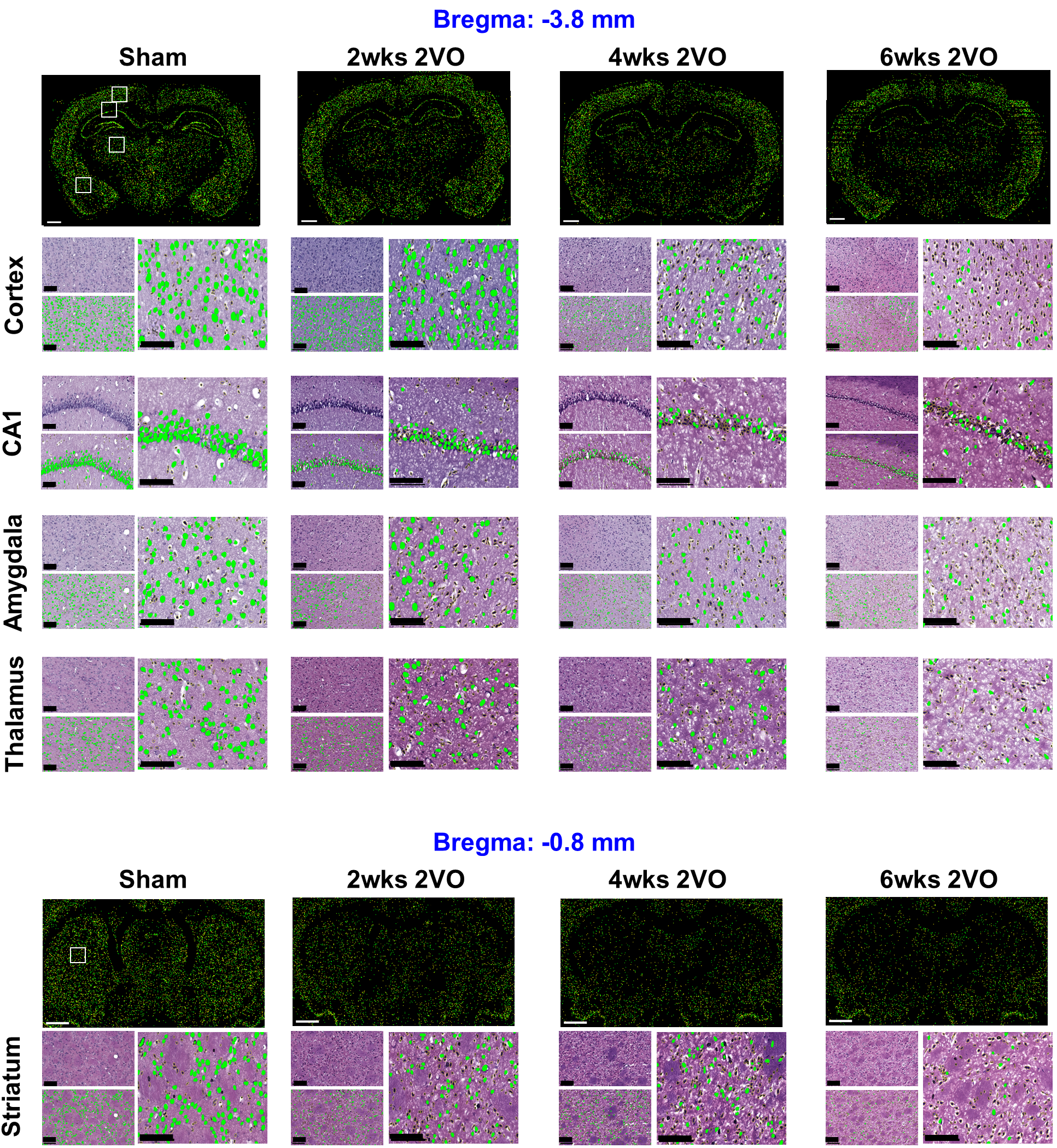

### Supplemental Figure 2

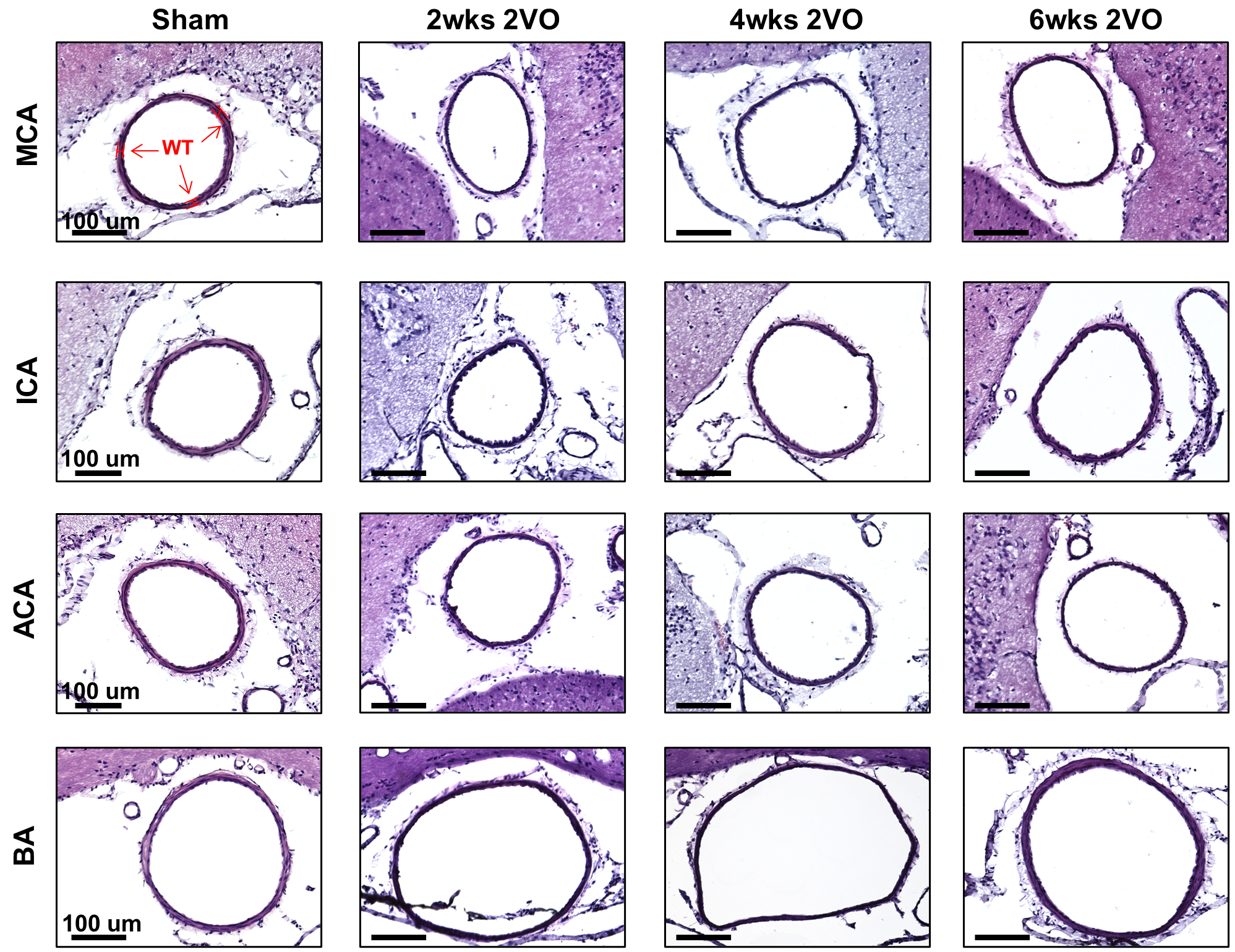

### Supplemental Figure 3

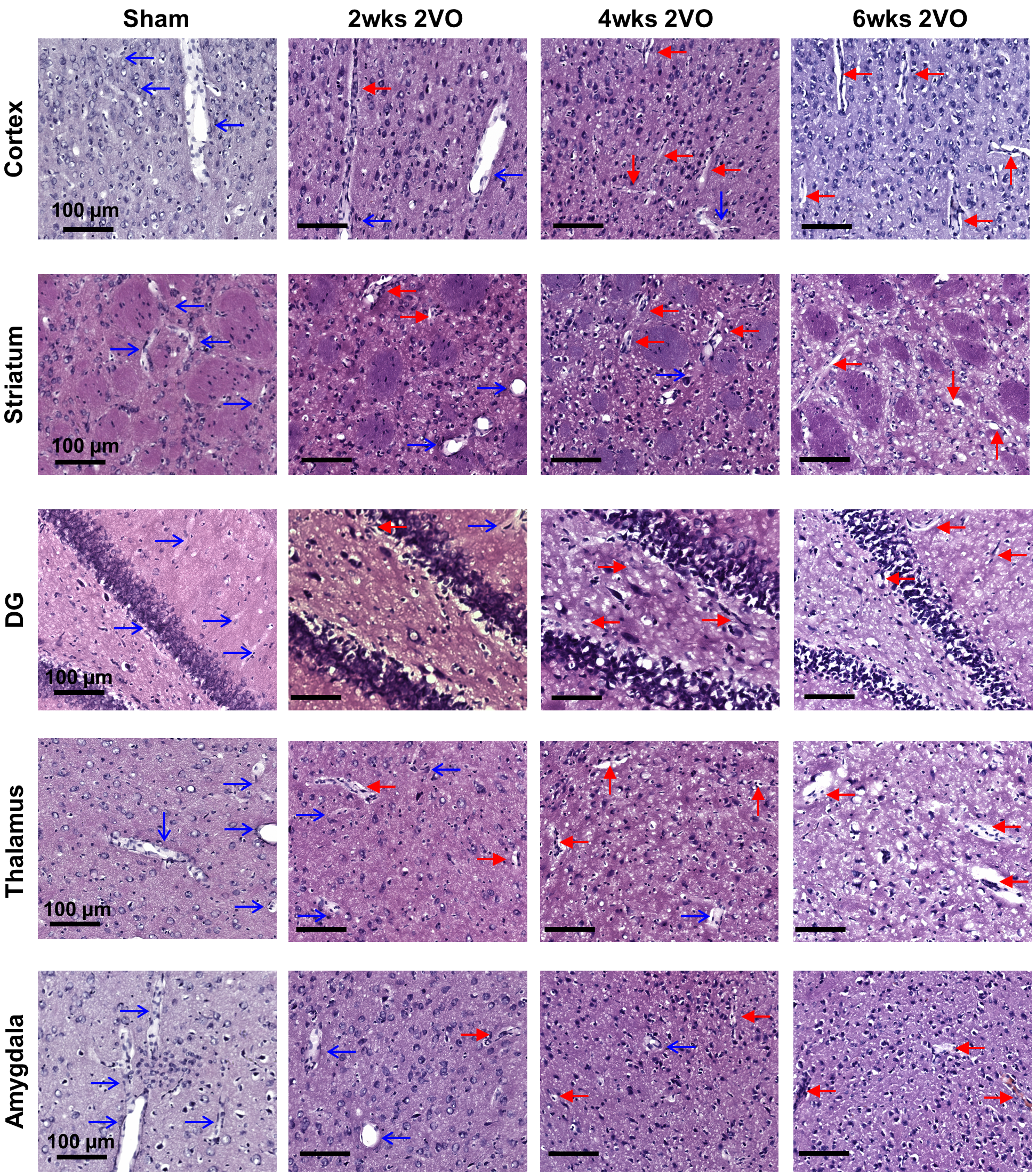

### Supplemental Figure 4

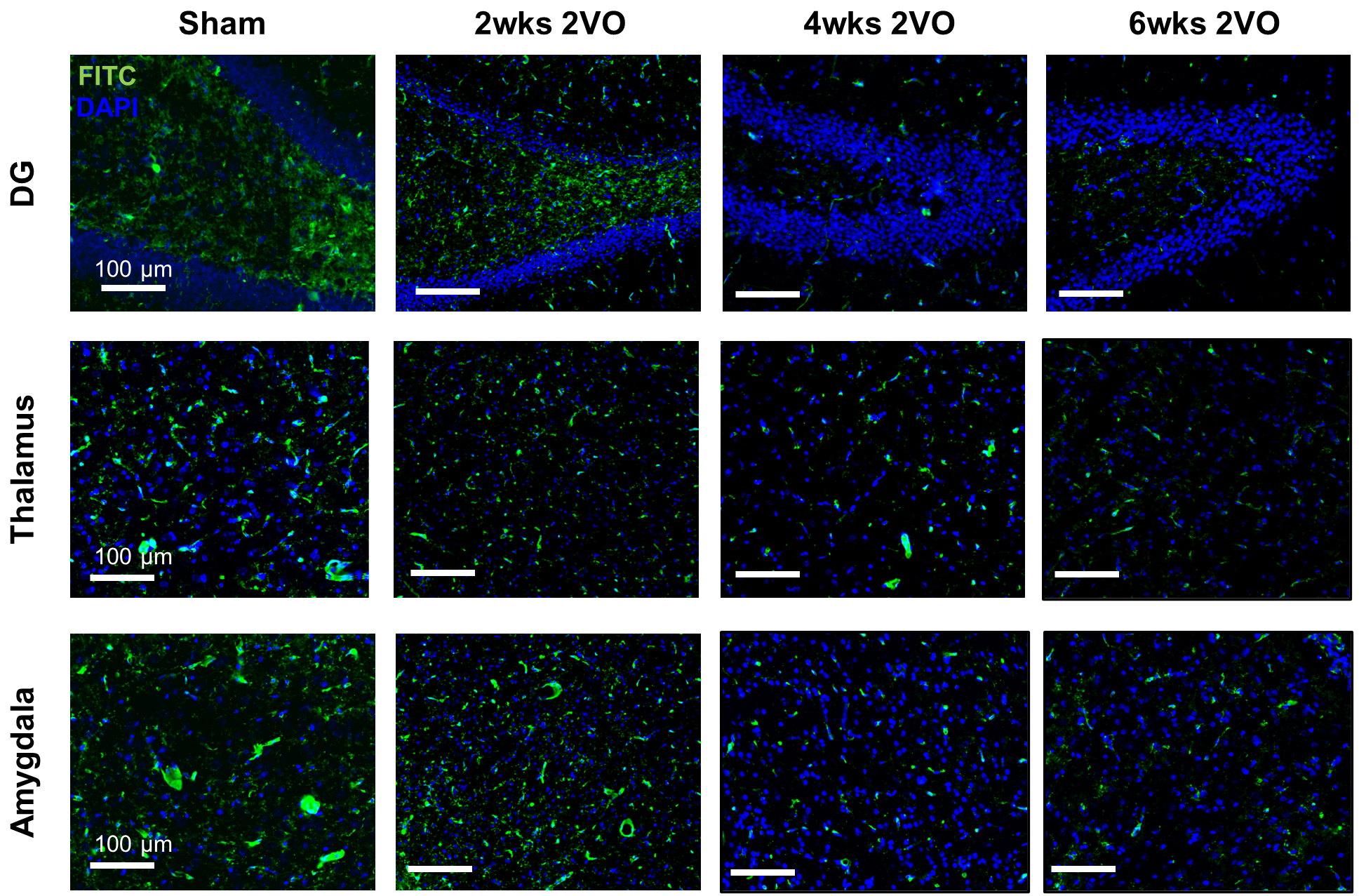

### Supplemental Figure 5

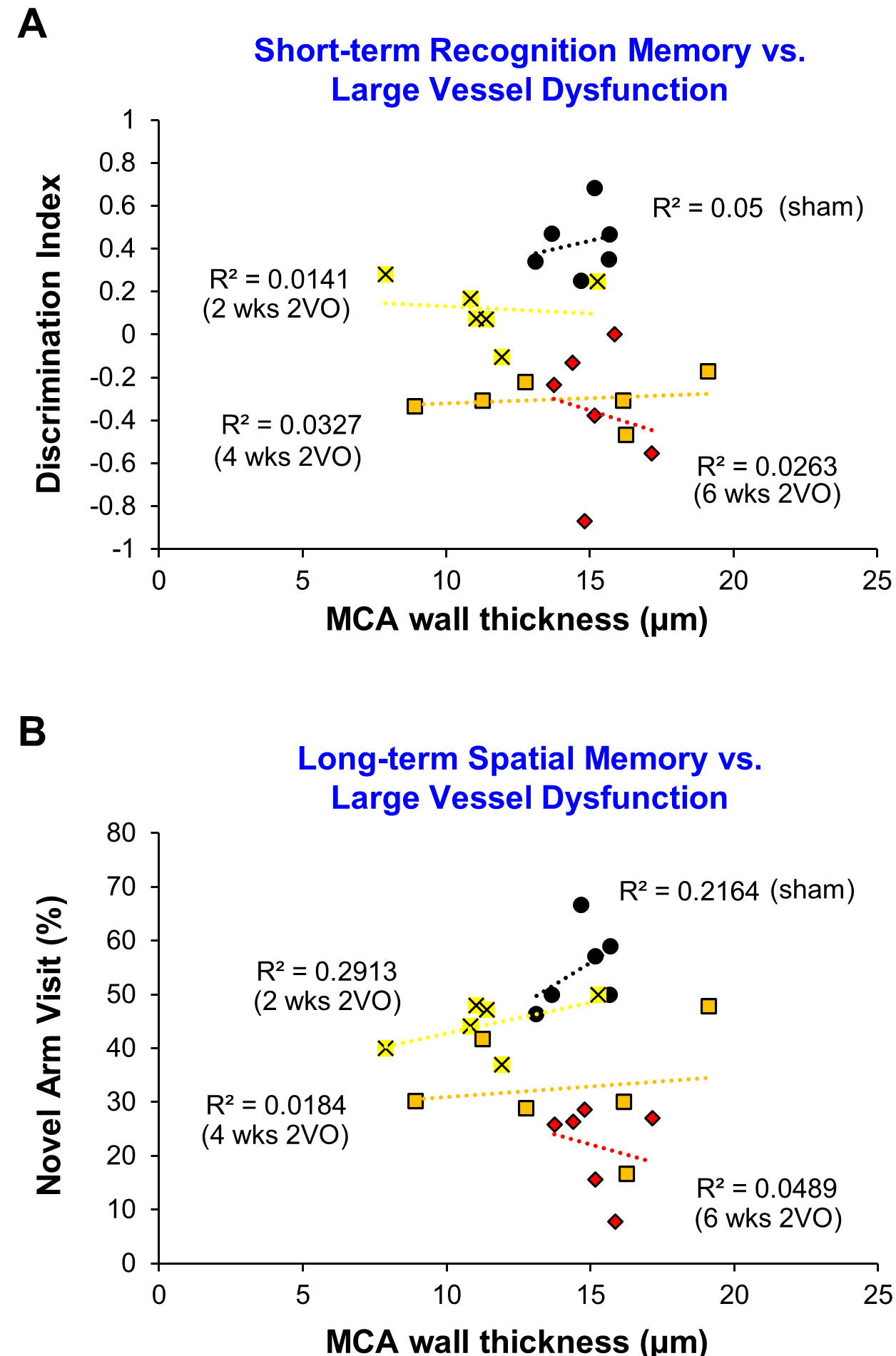
